# Stimulation-specific information is represented as local activity patterns across the brain

**DOI:** 10.1101/726414

**Authors:** Amirouche Sadoun, Tushar Chauhan, Samir Mameri, Yifan Zhang, Pascal Barone, Olivier Deguine, Kuzma Strelnikov

## Abstract

Modern neuroimaging represents three-dimensional brain activity, which varies across brain regions. It remains unknown whether activity within brain regions is organized in spatial configurations to reflect perceptual and cognitive processes. We developed a rotational cross-correlation method allowing a straightforward analysis of spatial activity patterns for the precise detection of the spatially correlated distributions of brain activity. Using several statistical approaches, we found that the seed patterns in the fusiform face area were robustly correlated to brain regions involved in face-specific representations. These regions differed from the non-specific visual network meaning that activity structure in the brain is locally preserved in stimulation-specific regions. Our findings indicate spatially correlated perceptual representations in cerebral activity and suggest that the 3D coding of the processed information is organized in locally preserved activity patterns. More generally, our results provide the first demonstration that information is represented and transmitted as local spatial configurations of brain activity.

## Introduction

Functional neuroimaging techniques represent the brain as a three-dimensional field of energy, which has a certain structure such as peaks, valleys etc. The classical analysis of brain activations focuses on finding the peaks of energy, but neglects the three-dimensional patterns of brain activity in the vicinity of the peaks. Several psychophysical and psychological studies have demonstrated that during perception, the brain processes spatial differences between perceived elements ^1^. E.g., this is evident in the visual system, where the differential approach begins in the retina, which consists of ON- and OFF-center receptive fields (see ^2^ for review). Due to the combination of excitatory and inhibitory connections, differences of activity between neuroglial populations are established as part of neural coding. This neural encoding, in turn, is the basis for low-level functions such as orientation detection ^3,4^, which are usually performed by clusters of cells with similar preferences (e.g. orientation columns). The spatial pattern formation is also seen in the primary visual cortex where some cortical layers may inhibit activity in the other layers ^5^. Though there is no doubt about the importance of spatial activity patterns, traditional fMRI connectivity studies employ correlation analyses across time-courses of individual voxels, which does not take into account any spatial correlations that might exist among the activity patterns. It is crucial to recognise that the brain is a three-dimensional structure which may be considered as an ensemble of small volumes called voxels, with the activity of each voxel representing a summary level of neuroglial activity ^6,7^. Therefore, in addition to a temporal component, neural codes may also have a strong spatial component observable as patterns of 3D activity ^8^. This possibility is also reinforced by the finding that repeating patterns of network information have been shown to account for up to 50% of the variance in fMRI blood oxygen level dependent (BOLD) data ^9,10^. Furthermore, Majeed et al. (2009) have also suggested that spatial patterns of activity could be explained by propagating wave of synchronized activity along the rat cortex. This raises the hypothesis that spatial configurations (patterns) of activity, which correspond to spatial differences of activity levels between the neighbouring neuroglial populations during a given stimulation, constitute an important aspect of information coding not only at the sensory stages, but also in higher cortical areas. An analysis of spatial patterns in fMRI data raises two questions: whether fMRI, as a technique, is sensitive enough to make meaningful measurements of local spatial activity patterns ^11^; and, if this is the case, how does one quantify correlations between local patterns.

Using gradient and divergence calculations we demonstrated ^12^ that differences of activity between adjacent voxels in certain loci of the brain are significant at the group level and are stimulation-dependent. To explain the inter-subject stability of the differential code, we suggested that the stimulation-related differences of activity between adjacent voxels in the BOLD signal contrast images are consistent. This, in our view, further indicates that there may be a spatial organisation of this information in the form of patterns of activity in the brain. Another proof for the existence of spatially coded information in the brain can be found in the results of multivoxel pattern analysis (MVPA), which makes use of the spatial differences in activation between voxels (see ^13,14^ for review). Despite its proven ability to classify brain activity patterns based on the stimulation, it lacks neurophysiological interpretation and has poor localizing ability ^15^. However, local approaches with MVPA are also possible, e g. in some cases it is possible to distinguish complex sounds on the basis of the auditory activity patterns ^16^. A more precise approach to the estimation of spatial distribution of brain activity would be to correlate a certain multivoxel spatial pattern in a region of interest with other spatial patterns in the brain, in other words, a localised cross-correlation analysis. The cross-correlation technique provides a unique correlation value per voxel, which summarizes the resemblance of spatial activity in the vicinity of this voxel with the seed pattern.

In this article, we present one possible implementation of such a cross-correlation analysis which allows us to estimate to what extent the variations of activity between the groups of voxels are similar in different brain regions. The existence of such correlated patterns of voxel activity could indicate local spatial coding in the brain. In line with our hypothesis, we further ask whether this local spatial coding in brain activity is specific to cognitive processing, and whether a given spatial pattern of activity can recur throughout various specialised cortical areas of the brain.

## Results

### Cross-correlation analysis in the occipital area

In order to test the specificity of our method we took a seed pattern in the fusiform face area of the face-specific contrast, coordinates: x = 33, y = −76, z = −8 mm, equivalent to x =16, y = 13, z = 15 voxels (here and in other places, we also indicated the voxel coordinates of the seed because they are used in the cross correlation toolbox). As expected, the most significant peak in the group analysis was found at the location which corresponds to the coordinates of the seed pattern (Tables 1 and 5) (a 1-2 voxel displacement is sometimes observed at the group level analysis, though exactly the same seed coordinates are found at the individual level).

The rest of the results section will omit the predictable correlations of 1 found at the seed location. The other significant peaks were found in the contralateral FFA, the right cerebellum, and the left orbitofrontal region (Fig. 1 a, b).

**Table 1.** Coordinates of significant correlations and their anatomic location. (Face vs. Random. Seed: right fusiform. x = 33, y = −76, z = −8 mm. Contrast: face-specific)

| Anatomic location | p value | Cluster size ($K_E$) | z value | x | y | z |
| --- | --- | --- | --- | --- | --- | --- |
| **R fusiform** | 0.0001* | 542(61) | > 6 | 33 | -76 | -8 |
| **R cerebellum** |  |  | 5.83 | 51 | -58 | -32 |
| **L fusiform** | 0.0001* | 123(1) | 5.50 | -30 | -79 | -8 |
| **L fusiform** | 0.036** | 16 | 4.34 | -42 | -49 | -17 |
| **L orbitofrontal** | 0.0001* | 51 | 4.71 | -12 | 44 | -17 |

\*FWE cluster corrected p-value. \*\*The FDR cluster corrected p-value. Cluster sizes are reported at the $p < 0.0001$ uncorrected level. In brackets cluster sizes are indicated at the FWE corrected level when appropriate.

**Table 2.** Coordinates of significant correlations and their anatomic location. (Face vs. Random. Seed: Left fusiform. x = −30, y = −79, z = −8 mm. Contrast: face-specific)

| Anatomic location | p value | Cluster size ($K_E$) | z value | x | y | z |
| --- | --- | --- | --- | --- | --- | --- |
| L fusiform | 0.0001* | 408(66) | > 6 | -30 | -79 | -8 |
| L fusiform |  |  | 5.38 | -36 | -49 | -14 |
| L cerebellum |  |  | 4.81 | -33 | -55 | -35 |
| R fusiform | 0.0001* | 336(1) | 5.15 | 36 | -40 | -17 |
| R inf. Temporal |  |  | 5.07 | 51 | -49 | -14 |
| R fusiform/inf. Occip. |  |  | 4.91 | 39 | -73 | -8 |

\*FWE cluster corrected p-value. Cluster sizes are reported at the $p < 0.0001$ uncorrected level. In brackets cluster sizes are indicated at the FWE corrected level when appropriate.

**Table 3.** The values of the cross-correlation for each subject in the fusiform areas (Contrast: face-specific, seed in the left fusiform area)

| Contrast | Face-specific |  |  |
| --- | --- | --- | --- |
| Location | Seed (Left Fusiform) | Fusiform Contralateral | Fusiform Contralateral |
| Coordinates<br>x y z | -30 -79 -8 | 39 -73 -8 | 33 -76 -8 |
| Subjects |  |  |  |
| 1 | 1 | 0.47 | 0.56 |
| 2 | 1 | 0.71 | 0.73 |
| 3 | 1 | 0.59 | 0.71 |
| 4 | 0.99 | 0.58 | 0.65 |
| 5 | 1 | 0.55 | 0.60 |
| 6 | 1 | 0.72 | 0.74 |
| 7 | 1 | 0.74 | 0.68 |
| 8 | 0.99 | 0.59 | 0.50 |
| 9 | 1 | 0.64 | 0.52 |
| 10 | 1 | 0.55 | 0.52 |
| 11 | 1 | 0.65 | 0.57 |
| 12 | 1 | 0.70 | 0.70 |
| 13 | 1 | 0.71 | 0.70 |
| 14 | 1 | 0.61 | 0.70 |
| 15 | 1 | 0.58 | 0.35 |
| 16 | 1 | 0.68 | 0.66 |
| **Mean** | **0.99** | **0.63** | **0.62** |
| **Bootstrap CI [inf, sup]** | **[0.99, 1]** | **[0.59, 0.66]** | **[0.56, 0.66]** |

Each subject's value indicates the result of the cross-correlation at the above mentioned coordinates. Mean values and bootstrap confidence intervals are indicated (CI; re-sampling 10000 times with a bias corrected and accelerated (BCa) percentile algorithm for confidence intervals<sup>50</sup>).

**Table 4.** Coordinates of significant correlations and their anatomic location. (Visual vs Random. Seed: right fusiform. x = 33, y = −76, z = −8 mm. Contrast: non-face-specific)

| Anatomic location | p (corr) Cluster level | Cluster size (K<sub>E</sub>) | z value | x | y | z |
| --- | --- | --- | --- | --- | --- | --- |
| **R fusiform** | 0.0001* | 1259(99) | > 7 | 33 | -73 | -8 |
| **R cerebellum** |  |  | 6.74 | 45 | -58 | -32 |
| **R lingual** |  |  | 6.62 | 24 | -88 | -5 |
| **L cerebellum** | 0.0001* | 2086(41) | 6.43 | -24 | -58 | -29 |
| **L lingual/ fusiform** |  |  | 6.37 | -18 | -88 | -8 |
| **R postcentral** |  |  | 5.90 | 48 | -25 | 43 |
| **L MCC (Middle Cingulate Cortex)** | 0.0001* | 557(6) | 5.82 | -9 | -4 | 46 |
| **L MCC** |  |  | 5.75 | 0 | -1 | 46 |

\*FWE cluster corrected p-value. Cluster sizes are reported at the $p < 0.0001$ uncorrected level. In brackets cluster sizes are indicated at the FWE corrected level when appropriate. Cognitive interpretations are limited for this contrast (non-specific visual activations).

**Table 5.** The values of cross-correlation for each subject in the fusiform, visual and other areas (Contrasts: face-specific and non-face specific contrasts; seed in the right fusiform area)

| Contrast | face-specific |  |  |  | non-face specific |  |  |  |
| --- | --- | --- | --- | --- | --- | --- | --- | --- |
| Location | Seed (right fusiform) | Fusiform Contralateral | L Orbitofrontal | R Cerebellum | Seed (right fusiform) | Seed vicinity | Occipital (L Lingual/ Fusiform) | R lingual |
| Coordinates<br>x y z | 33 -76 -8 | -30 -79 -8 | -12 44 -17 | 51 -58 -32 | 33 -76 -8 | 33 -73 -8 | -18 -88 -8 | 24 -88 -5 |
| Subjects |  |  |  |  |  |  |  |  |
| 1 | 0.99 | 0.61 | 0.71 | 0.67 | 0.99 | 0.93 | 0.74 | 0.76 |
| 2 | 1 | 0.83 | 0.71 | 0.77 | 1 | 0.96 | 0.79 | 0.78 |
| 3 | 1 | 0.66 | 0.38 | 0.68 | 1 | 0.95 | 0.74 | 0.58 |
| 4 | 1 | 0.76 | 0.74 | 0.80 | 1 | 0.92 | 0.70 | 0.64 |
| 5 | 1 | 0.64 | 0.69 | 0.75 | 0.99 | 0.93 | 0.72 | 0.69 |
| 6 | 1 | 0.65 | 0.59 | 0.63 | 1 | 0.91 | 0.70 | 0.73 |
| 7 | 1 | 0.72 | 0.66 | 0.55 | 1 | 0.92 | 0.74 | 0.73 |
| 8 | 1 | 0.54 | 0.50 | 0.76 | 0.99 | 0.94 | 0.77 | 0.77 |
| 9 | 1 | 0.58 | 0.49 | 0.62 | 0.99 | 0.94 | 0.78 | 0.73 |
| 10 | 1 | 0.53 | 0.63 | 0.55 | 0.99 | 0.87 | 0.77 | 0.74 |
| 11 | 1 | 0.62 | 0.69 | 0.83 | 1 | 0.92 | 0.69 | 0.68 |
| 12 | 0.99 | 0.69 | 0.54 | 0.65 | 0.99 | 0.93 | 0.78 | 0.74 |
| 13 | 1 | 0.75 | 0.75 | 0.84 | 1 | 0.95 | 0.80 | 0.72 |
| 14 | 1 | 0.64 | 0.72 | 0.77 | 1 | 0.95 | 0.80 | 0.69 |
| 15 | 1 | 0.52 | 0.71 | 0.76 | 0.99 | 0.89 | 0.81 | 0.78 |
| 16 | 1 | 0.76 | 0.66 | 0.76 | 1 | 0.95 | 0.79 | 0.77 |
| **Mean** | **0.99** | **0.66** | **0.64** | **0.71** | **0.99** | **0.93** | **0.76** | **0.72** |
| **Bootstrap CI [inf, sup]** | **[0.99, 1]** | **[0.61, 0.70]** | **[0.57, 0.68]** | **[0.67, 0.75]** | **[0.99, 1]** | **[0.92, 0.94]** | **[0.74, 0.77]** | **[0.69, 0.74]** |

Each subject's value indicates the result of the cross-correlation in the above-mentioned coordinates. Mean values and bootstrap confidence intervals (BCa, $p < 0.05$ ) are indicated.

**Fig. 1.**
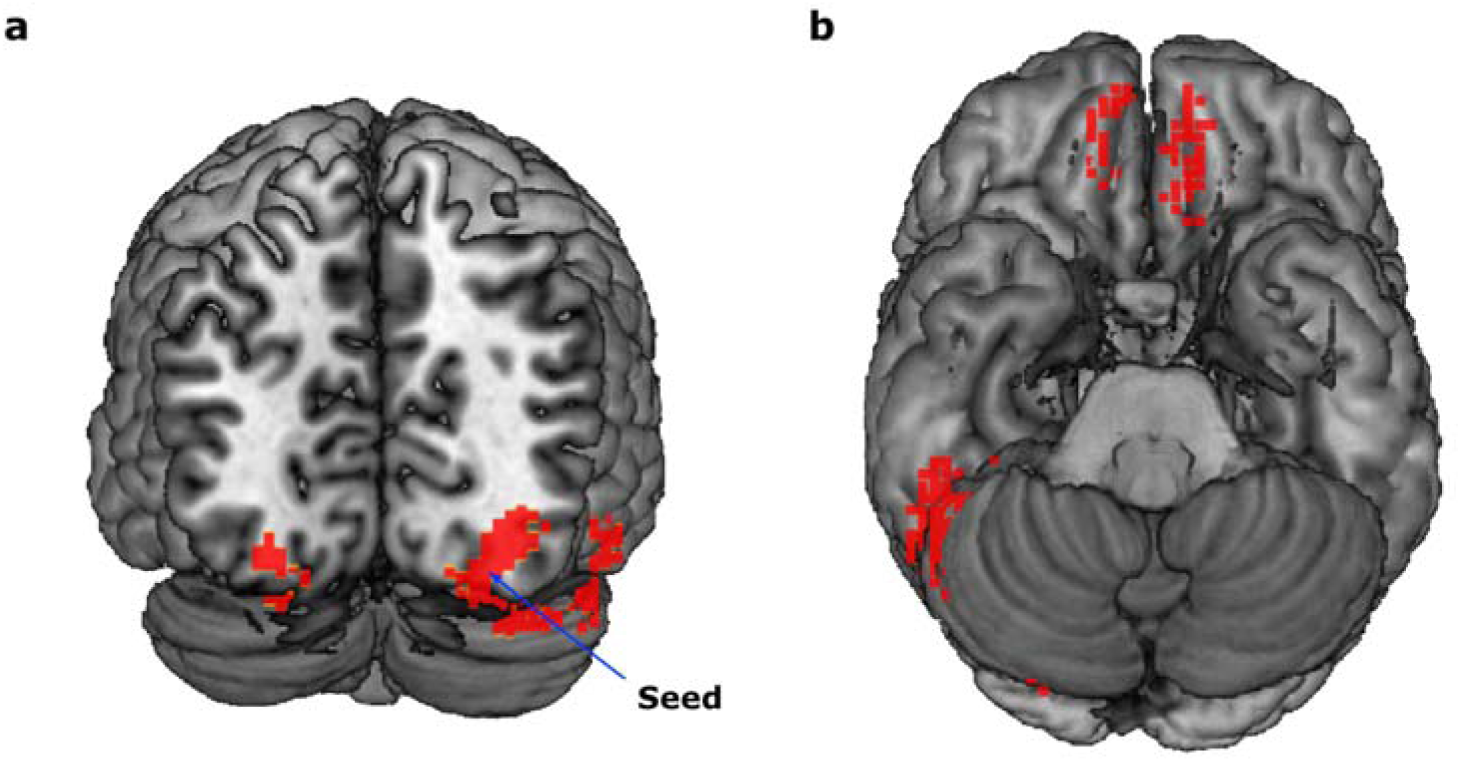
Significant clusters resulting from the cross-correlation analysis in the face-specific contrast. **a**, Ipsi and contralateral significant clusters are shown in the occipital lobes (FFA) and the cerebellum on the 3D template map. **b**, Significant clusters in the orbitofrontal areas. In both images, clusters are shown at the uncorrected threshold for illustrative purposes. The arrow indicates the seed. MRICron software was used to display the figures (https://www.nitrc.org/projects/mricron).

To verify the consistency of the analysis, we used the contralateral peak from the previous analysis as the seed pattern. If the technique is robust this reverse-seed analysis should be able to detect the initial seed. In fact, the most significant correlation peaks of this reverse-seed analysis were indeed found at the original seed and its vicinity (Fig. 2a; Tables 2 and 3). We further verified that this reverse-seed correlation holds for each subject (mean = 0.62, bootstrapped CI = [0.55, 0.66]). Thus, a pattern in the opposite hemisphere can be detected whatever the side of the seed pattern used for the analysis.

**Fig. 2.**
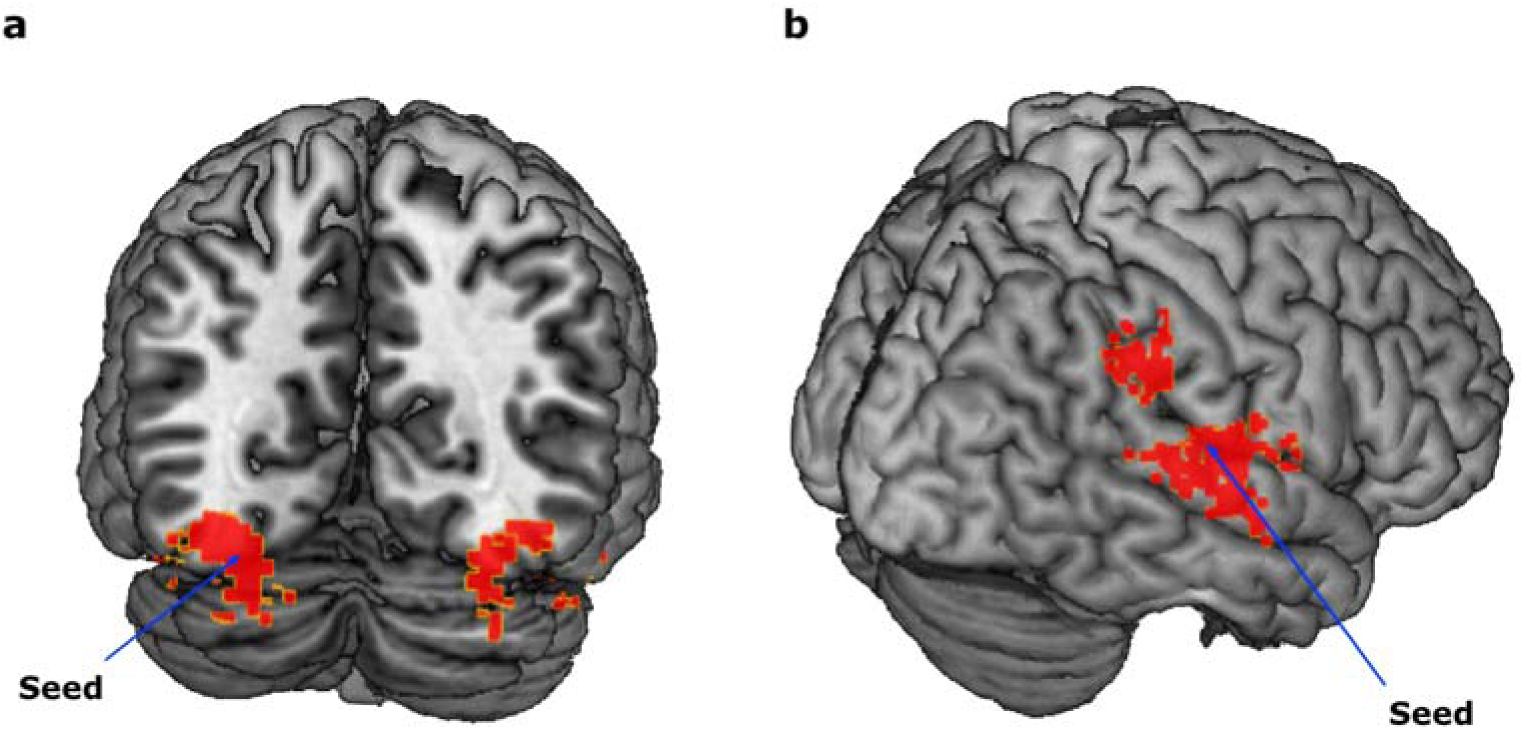
Significant clusters with the seed pattern in the contralateral FFA or in the primary auditory area. **a**, Significant clusters in the FFA resulting from cross-correlation in the face-specific contrast with the seed on the contralateral FFA. Ipsi and contralateral significant correlations are shown on the 3D template map. **b**, Significant clusters in the auditory area resulting from cross-correlation in the face-specific contrast. In both images, clusters are shown at the uncorrected threshold for illustrative purposes. Arrows indicate the seed. MRICron software was used to display the figures (https://www.nitrc.org/projects/mricron).

Next, to verify, as a control, if the method provides other results for non-face-specific data, we analyzed the non-face-specific contrast with the seed pattern at the same coordinates as for the above described face-specific contrast (x = 33, y = − 76 and z = − 8 mm ⇔ x = 16, y = 13, z = 15 voxels)(Table 4).

In addition to the seed coordinates, we found significant peaks in the occipital region (ex: x = 24, y = −88, z = −5 mm; p < 0.001, FWE correction). These peaks did not coincide with those found in the face-specific contrast.

Thus, significant correlations with a seed pattern can be found across the hemispheres both for face-specific and non-face-specific contrasts but at different locations. Furthermore, values of the cross-correlations per subject were above 0.5 and the mean correlation values for non-seed areas were in the range 0.6-0.8 (Table 5).

Likewise, we noticed that at the seed coordinates the values of the cross-correlation are 1 or 0.999 for all the subjects. This indicates the high specificity and accuracy of our method.

Consideration of the rotation angles for each significant peak in the cross-correlation results showed that the rotated seed pattern at the coordinates of the maximum significant correlation was approximately tangential to the cortex at these coordinates (for example, a seed pattern in the occipital region was rotated almost by 90 degrees for a correlation peak in the orbitofrontal region). This reflects the natural-occurring relative geometry between brain regions.

Non-parametric analysis with SnPM toolbox gave similar results indicating the robustness of our analysis (see supplementary information).

### Cross-correlation analysis with auditory activity

To verify the specificity of our method with respect to sensory modalities, we used the seed patterns defined in the auditory cortex for cross-correlation with both face-specific and non-specific contrasts (at the coordinates x = 10, y = 32, z = 21 voxels ⇔ x = 51, y = −19, z= 10 mm). Here, the seed pattern is outside the stimulation specific areas. The most significant correlation peaks were found at the seed pattern and its vicinity of the auditory cortex (see supplementary Table S1). This suggests that our technique is stimulation specific.

Conversely, we also compared the images resulting from a seed pattern in the fusiform face area with images resulting from the seed pattern in the auditory region (which served in this case as a non-specific statistical baseline). The results of this analysis (Table 6) were similar to those obtained using randomised images as the statistical baseline (see Methods for details).

**Table 6.** Coordinates of significant correlations using cross-correlation with the auditory seed as statistical baseline (Face-specific contrast).

| Anatomic location | p value | Cluster size ($K_E$) | z value | x | y | z |
| --- | --- | --- | --- | --- | --- | --- |
| **R fusiform** | 0.0001* | 944(80) | > 7 | 33 | -76 | -8 |
| **R cerebellum** |  |  | 6.27 | 51 | -49 | -35 |
| **L fusiform** | 0.0001* | 434(5) | 5.32 | -24 | -82 | -11 |
| **L orbitofrontal** | 0.001* | 230 | 4.84 | -12 | 47 | -8 |

\*FWE cluster corrected p-value. Cluster sizes are reported at the $p < 0.0001$ uncorrected level. In brackets cluster sizes are indicated at the FWE corrected level when appropriate.

## Discussion

### Face processing viewed through long-range spatial correlations

In the present work, we propose a new approach to analyze fMRI data using spatial correlations. As a proof-of-concept, we apply our approach to a publicly available fMRI dataset (see Methods), and show that our methodology allows us to find specific spatial patterns of the activity following the presentation of facial visual stimuli. These patterns are transmitted to the contralateral FFA with high correlation in both group and individual level analysis, indicating the conservation of spatial information regarding faces in both left and right FFA. In addition, by performing reverse-seed correlations (setting the seed in the contralateral area) we found the same results in the FFA – confirming both robustness and a high specificity of our approach. In our results, we found that the correlations are high (up to 0.8), suggesting that a large amount of spatial information in brain activity is transferred between these areas.

The FFA is involved in face perception, and is known to respond actively to face stimuli ^17–21^. However, the spatial conservation and transmission of activity-related information has never been observed. The present work provides the first demonstration of stable spatial patterns of brain activity in response to face stimuli, which preserve information across brain structures. In addition, we also find a high correlation in the orbitofrontal cortex, which is in line with several findings which have demonstrated the existence of face-selective neurons in the orbitofrontal cortex ^20,22,23^, and shown that this region is actively involved in face recognition ^20,22–25^.

Barat and collaborators (2018) showed that orbitofrontal face cells encode facial stimuli by, first, discriminating them from non-facial ones, and thereafter categorizing them according to their social and emotional dimensions. Face neurons encode these aspects despite differences in face stimuli concerning, for example, identity or head position of the face ^22^. In the present study, we used the dataset from a work where the presented face-stimuli have various expressions (generally, happy or neutral). The presence of significant correlation in the orbitofrontal cortex in our findings indicate that the social and emotional dimensions that are carried by faces are represented as spatial patterns of activity. Similarly, we also found significant cerebellar correlations for face-stimuli – an observation that is in agreement with studies which have implicated the cerebellum in facial information analysis ^20,26–28^.

Thus, the obtained correlations are highly specific to facial information. They are found in the FFA, and the orbitofrontal and cerebellum regions, which are known for being activated after facial stimuli presentations in several studies as described above. In addition to the above agreement with literature, another argument for the stimulation specificity of our results is the fact that the spatial correlation analysis was performed on the “stimulation vs. baseline” contrasts and all the images were acquired under the same conditions and processed in the same way excluding acquisition and processing artefacts in contrast images. In addition, we also showed that using a seed in the auditory region produces high correlations in the auditory areas – further indication of the specificity of our methodology.

### General considerations of local activity patterns

Our analysis shows that correlated stimulation specific patterns are found throughout the brain. Moreover, this suggests that there are recurring local patterns of meaningful stimulus-related activity. One interpretation of these local patterns could be that there is a partial conservation of stimulus-related information in localised regions in the cortex, and that the activity of any single unit is related to the activity of the neighbouring units. The maintenance of these local patterns would also be energetically favourable as compared to longer range activity patterns, and could be an efficient neuro-glial code for local copies of stimulus-related information. Likewise, we suppose that brain information is spatially organized in such a way that each neuro-glial population, represented by a given voxel, maintains a given level of activation according to its neighbours to maintain this pattern stable. Maintaining patterns stable is also probably related to the necessity to spend less energy in an optimal way to maintain information-representation in close vicinity instead of using the long-distance arrangements.

Spatial analysis is widely spread in electrophysiological research, e.g., to analyze the structure of cortical activity; spatial differentiation is widely used in electrophysiological studies in the form of current source density (CSD) analysis ^29,30^. In the latter, spatial differentiation permits to calculate electric current flows, given the measured electric potentials. Though the BOLD signal is an indirect and complex measure of brain activity ^31^, its correlation with electrical measurements in the brain has been established by numerous studies (e.g., ^32–36^. Given that electromagnetic and metabolic energies are highly correlated during brain activity, Friston ^37^ proposed that the most suitable summary form of energy to describe brain mechanisms is free energy. Physically, free energy represents a difference between internal energy and the product of entropy and temperature. In the case of stable temperature, the most important parameter that influences free energy will be entropy. In our previous research, we investigated the possibility to apply mathematical formalism, which is similar to the electrophysiological current source density (CSD) analysis ^29,30^, to the BOLD signal contrast maps. In the biophysical consideration of this analysis ^12^, we indicated that in line with the free energy minimization principle ^38^, gradients of energy between voxels should spontaneously disappear with time. However, we observed stable task-related gradients of activity at the group level ^12^, necessitating the existence of the stimulation-related processes which act to maintain the described spatial gradients. These oxygenation related gradients are likely to be driven by electrical gradients which encode the flow of local information. The results from our spatial-correlation analysis are in favour of this interpretation. We compared fMRI, EEG and MEG spatial differential activity during different tasks to see what amount of fMRI differential activity corresponds to the electromagnetic differential activity ^39^. Distributed source reconstruction was used to obtain 3-dimensional models of electric and magnetic activity in EEG and MEG prior to spatial differentiation. Using independent datasets with the same stimulation, we demonstrated that the mean spatial overlap of the fMRI differential activity with EEG and MEG may be about 80%. In addition, about 93% of divergence (spatial sources) in fMRI corresponded to the EEG and MEG divergence. Furthermore, previous studies have shown that glutamate related excitatory synaptic transmission accounts for about 70% of total energy turnover ^6^ in the brain, whereas GABAergic processes account for only about 15% of total energy turnover by neurons and glia ^40^ – thus suggesting that the stable local BOLD patterns identified in our analysis are results of local excitatory interactions between neurons.

Since brain activity is dynamic in time, it is important to understand how average BOLD contrasts can lead to local patterns of activity. To account for the stability in time of the observed energy flows and resulting spatial patterns, one can suggest that the repetitive stimulation-related patterns of energy flows in the brain form the average patterns reflected in the BOLD signal contrasted images. This is in line with what we find in our results, where spread of energy flow appears to be conserved and stable in a 3D space, as indicated by patterns of activation. Information-related activity patterns are closely related to the spatial distribution of stable energy flows in the brain. Energy flows in the brain are generally defined as coherent spatial and temporal changes in the energy turnover of neuroglial units related to information treatment ^7^; these flows are the result of the stimulation-driven transformations of energy that propagate in certain directions along the cellular structures (axons, dendrites, synapses, etc.) in neuroglial networks. When a neural signal reaches a neuroglial population, it increases the internal energy in this neuroglial population. These connections can be detectable only at their arrival point when they cause an abrupt increase of activity in the specialized population. The above described link between the directions of gradients and energy flows was confirmed by the finding that the sources, from which energy flows spread in the cortex, were in the occipital cortex during face processing and in the superior temporal cortex during auditory word processing ^12^. Patterns of brain activity may be related to integrative cognitive processes ^41^.

We think that future studies using spatial cross-correlation analysis combined with MVPA could be very informative about whether activity-based decoding methods could be more efficient in the regions where correlated spatial patterns are observed. The spatial connectivity approach may complement the data on intrinsic connectivity of homotopic brain areas ^42^. A fundamentally important question is whether during multisensory interactions patterns of brain activity carry the similar spatial information across modalities, these studies can further develop the models of such phenomena as McGurk effect ^43^. Moreover, in pathologies such as multiple sclerosis, epilepsy and Alzheimer disease spatial cross correlations could be a useful methodology for testing how patterns of structure and activity are distributed throughout the cortex ^44^ in addition to identifying how these patterns of activity may change in early stages of these diseases, and also during the follow up of the patients, leading to a powerful new diagnosis tool, in addition to the existing techniques ^45–47^.

## Materials and methods

### fMRI dataset

In order to analyze functional pattern similarities in the brain during a given task, the freely available data were chosen from a work of Wakeman and Henson ^48^ (ftp://ftp.mrc-cbu.cam.ac.uk/personal/rik.henson/wakemandg_hensonrn/). The Matlab scripts attached to the original dataset were used for pre-processing and statistical analysis (using the Matlab SPM toolbox), resulting in contrast images in the MNI space with 3mm isotropic voxels smoothed by a 3D 8mm isotropic Gaussian kernel. In the original study, the subjects (n = 16) were presented gray scale images of familiar and unfamiliar faces, and faces scrambled by a 2-D Fourier transform. Original and scrambled faces were cropped using a mask built on the basis of a combination of familiar and unfamiliar faces.

### Toolbox description

We built a toolbox in C++ using ITK (Insight Toolkit (https://itk.org/)) allowing cross-correlation analysis using the Fourier transform of the 3D patterns. The seed 3D pattern taken at a given location in the brain was rotated in steps of 10 degrees along the z, y and x axes, and for each rotation step the pattern was cross-correlated with the entire brain activity (Fig. 3a and b). Given 18 steps of rotations around each axis to obtain all possible non-redundant rotations, the total number of configurations was 18×18×18 = 5832. Since both the rotations and the cross-correlation analysis are computationally intensive, it is important to accelerate the analysis using Fourier transform. The analysis of one subject took about 43 minutes on the PC optimized for performance (available in Windows options). The whole analysis on 16 subjects took about 4 hours. The maximal value of correlation for each rotation step was saved as a voxel value in the NIFTI file alongside with NIFTI files for the x, y and z angles of rotation corresponding to this correlation. The main executable file in our toolbox (cross.exe) takes as input the name of the NIFTI file and the coordinates of the seed pattern (tested with Windows 10).

**Fig. 3.**
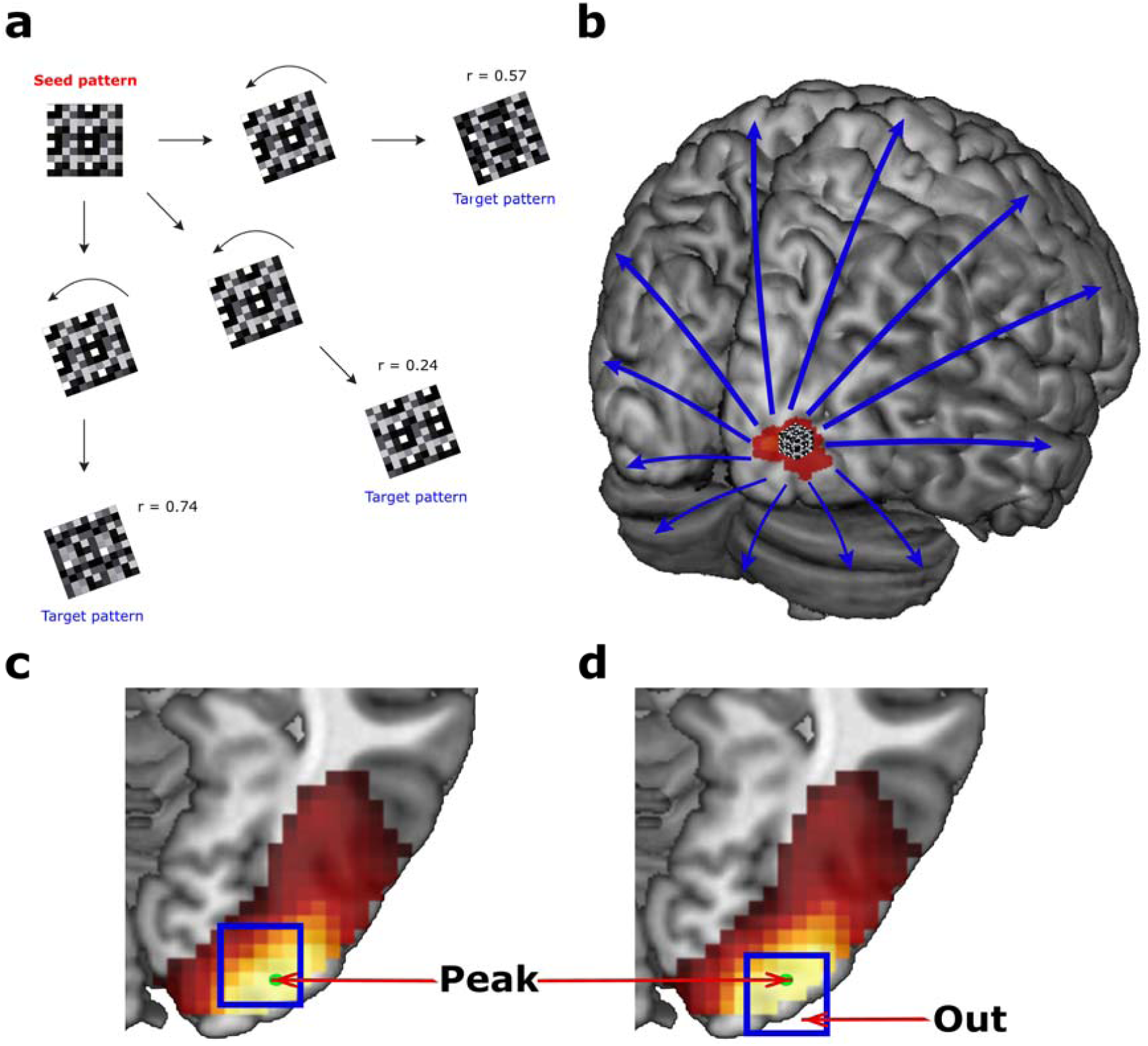
Schematic representation of the procedure of cross correlation. **a**, The seed pattern is turned then compared with target patterns. **b**, Activity in the seed pattern is cross-correlated with the entire brain activity. **c**, An example of a peak of activity where the pattern is centered deeper in the brain with respect to the peak. **d**, An example of a pattern that has a large part outside of the brain (indicated by the word ‘Out’ with arrow) – the situation to be avoided by moving the pattern deeper. In both **c** and **d**, the square represents the pattern and the dot inside the location of the peak of activity. MRICron software was used to display the figures (https://www.nitrc.org/projects/mricron).

### Data pre-processing

We put the obtained result (crosscorr_final.nii) (see supporting information) into the same MNI space as the original image using the 3D matrix in the intial SPM contrast image. In addition, given the presence of a small shift between the cross-correlation image and the initial contrast, we used a translation of the obtained result in Matlab (v 2017a). The obtained file was renamed (crosscorr_final_co_trans.nii). In order to verify if the latter has correctly been realigned to the initial contrast, we superimposed both images in SPM and verified that the peak in the cross-correlation image corresponds to the chosen coordinates in the initial contrast. Furthermore, we z-transformed the data after subtracting the mean to approximate the normal distribution. The resulting files were then renamed (Ztransf_Crosscorr_final_co_transl.nii). Thereafter, we created randomized brain images for every subject on the basis of the z-transformed images in order to use them as a reference for the statistical analysis. Indeed, in order to be considered statistically significant, a given data should be significantly different from a random distribution of the same type of data.

### Data analysis

SPM12 toolbox (https://www.fil.ion.ucl.ac.uk/spm/software/spm12/) was used to analyze the data. We used the two-sample t-test to compare the obtained z-transformed images and the randomized images (FWE correction, p < 0.05, extent threshold = 0). In order to ensure that there were no spurious correlations, we also used the two-sample t-test on the z-transformed images to compare the visual and auditory seed regions from the same initial contrast image.

Given the nature of the stimuli used by Wakeman and Henson, we chose two regions of interests in our analysis: an occipital and a temporal one. In the occipital region, we considered the coordinates corresponding to the peak of the brain activity in the contrasts. In the temporal one, as a control region, we choose the coordinates of the primary auditory area which should not show activity during visual stimulation. We used the contrasts con_0006.img (Faces (familiar + unfamiliar) > scrambled) and con_0005.img (scrambled > baseline) of every subject (the numbers of the contrasts are indicated according to the attached to the dataset Matlab scripts). We considered two seed points, the first one at the peak of the activity (Fig. 3c and d) in the two contrasts (x= 33, y = − 76, z = − 8 mm) in the Fusiform Face Area (FFA), (⇔ x = 16, y = 13, z = 15 voxels in matrix coordinates found from SPM Image view), the second peak (x = 51, y = −19, z = 10 mm) in the temporal region (⇔ x = 10, y = 32, z = 21 voxels in matrix coordinates, corresponding to the middle of the primary auditory area ^49^). The corresponding coordinates’ areas were verified in the SPM Anatomy (Version 2.2b) and xjView (http://www.alivelearn.net/xjview) toolboxes.

We used the matrix coordinates to perform the cross-correlation analysis with a radius of 5 voxels. When the activity peak is at the periphery of the brain, it is not useful to take the center of the pattern at the peak of the activity because in this case a large part of the pattern would be located outside the brain. When peaks are peripheral, it is better to center the pattern deeper into the brain with respect to the peak of activity so that the periphery of the pattern includes the peak (see Fig. 3c and d). We verified whether the considered pattern of activity with a radius of 5 voxels contains the peak of the activity. Given that the center of the pattern is the point *q*, the distance between that peak *p* and the point *q* should be inferior to the chosen radius for the pattern. In order to calculate that distance, we used the Euclidean distance *d(p,q)* between the peak of activity *p* and the point *q* in the chosen coordinates (1):

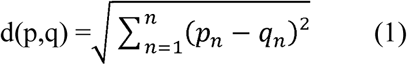

The obtained distance was 4.12 voxels, so inferior to the radius of 5 voxels.

To ensure the robustness of the cross-correlation analysis, we employed a seed-reversal approach. In this approach, the peaks of a given correlation-analysis were used as a new seed (in the contralateral FFA). If the technique is robust, this seed-reversal should lead us back to the initial seed.

Likewise, at the strongest peak of significance in the group analysis, we verified in each subject if the correlations at these coordinates are not different from 1 (Table 3).

In addition, given the distribution of the resulting data, which may not always respect the normal one, we also replicated the same analysis with a non-parametric approach using the SnPM toolbox (https://warwick.ac.uk/fac/sci/statistics/staff/academic-research/nichols/software/snpm) (two sample t-test, number of permutations = 256, variance smoothing = [8, 8, 8]). In this case, we did not apply the z-transformation to the images; we only put them into the same MNI space as the original contrast and performed the above-mentioned translation. This analysis is included in the supporting information part.

## Materials & Correspondence

Correspondence and requests for materials should be addressed to A.S. or K.S (, respectively).

## Data and scripts availability

Data generated and analyzed during the current study as well as the cross program and the MatLab scripts will be made available upon publication.

## Acknowledgements

We thank Daniel Wakeman and Richard Henson for data availability, data citation:

Wakeman, D. G. & Henson, R. N. OpenfMRI ds000117 (2014).

The authors are very grateful to the CerCo laboratory for supporting this work.

## Author Contributions

A.S. designed the study, analyzed the data, and prepared the manuscript. T. C., S. M., Y. Z., P.B., O.D. prepared the manuscript and discussed the results. K. S. designed the cross-correlation in ITK C++, designed the study, prepared the manuscript, and supervised the project. All authors reviewed the manuscript and discussed the results.

