## supporting information for "Stimulation-specific information is represented as local activity patterns across the brain"

### Supplementary results

#### Analysis with SPM

**Table S1 Coordinates of significant correlations and their anatomic location. (Auditory vs. Random. Seed: Right auditory area. x = 51, y = -19, z = 10 mm. Contrast: face specific)**

| **Anatomic location** | **p value** | **Cluster size (K_E_ )** | **z value** | **x** | **y** | **z** |
| --- | --- | --- | --- | --- | --- | --- |
| **R primary auditory area (Heschl)** | 0.0001* | 189(44) | > 6 | 51 | -19 | 10 |
| **R SupraMarginal** |  |  | 5.08 | 66 | -19 | 19 |
| **R sup. Temporal Gyrus** |  |  | 4.89 | 66 | -19 | 1 |
| **R SupraMarginal** | 0.0001* | 39 | 4.47 | 60 | -37 | 43 |
| **R MCC** | 0.005* | 28 | 4.42 | 6 | -13 | 37 |

*FWE cluster corrected p-value. Cluster sizes are reported at the p < 0.0001 uncorrected level. In brackets cluster sizes are indicated at the FWE corrected level when appropriate.

#### Analysis with SnPM

##### Cross-correlation analysis in the fusiform area

Similarly as previously with SPM analysis, we took a seed pattern in the fusiform area of the face-specific contrast (coordinates x =16, y = 13, z = 15 voxels ⬄ x = 33, y = -76, z = -8 mm). The most significant peak in the group analysis was found here at the same location, which corresponds to the coordinates of the seed pattern (x = 33, y = - 76 and z = - 8 mm, p = 0.003, FWE correction). As in SPM, we found significant correlations were in the contralateral fusiform area and the cerebellum regions (Table S2).

**Table S2 Coordinates of significant correlations and their anatomic location. (Face vs. Random. Seed: right fusiform. x = 33, y = -76, z = -8 mm. Contrast: con_0006)**

| **Anatomic location** | **p (corr) Voxel level** | **Cluster size (K)** | **Pseudo t** | **x** | **y** | **z** |
| --- | --- | --- | --- | --- | --- | --- |
| **R fusiform** | 0.0039* | 70 | 12.15 | 33 | -76 | -8 |
| **L fusiform** | 0.0039* | 1 | 7 | -24 | -79 | -11 |
| **R inf temporal** | 0.0039* | 8 | 6.21 | 57 | -55 | -23 |
| **R cerebellum** | 0.0039* | 25 | 6.03 | 51 | -58 | -35 |

*FWE corrected p-value. Cluster sizes are reported at the p = 0.0039 (permutations).

Regarding the contralateral occipital region, we took, as in the SPM analysis, the coordinates x = 37, y = 12, z = 15 vxls (⬄x = -30, y = -79, z = -8 mm). We found the most significant correlation peak was present at x= -30, y = -82 and z -8 mm (⬄ x =37, y = 11, z = 15 vxl, p = 0.0039; FWE correction) (Table S3) that was near the seed pattern coordinates. Verifying the significativity in the seed coordinates, we found it significant. In addition, significant correlations were found in the left and right fusiform areas and also in the cerebellum. Thus, similarly as in the SPM analysis, a pattern in the opposite hemisphere can be detected whatever the side of the seed pattern used for the analysis. This proves the stability of our method.

**Table S3 Coordinates of significant correlations and their anatomic location. (Face vs. Random. Seed: Left fusiform. x = -30, y = -79, z = -8 mm. Contrast: face specific)**

| **Anatomic location** | **p (corr) Voxel level** | **Cluster size (K)** | **Pseudo t** | **x** | **y** | **z** |
| --- | --- | --- | --- | --- | --- | --- |
| **L inf. Occipital/ L Fusiform** | 0.0039* | 90 | 11.66 | -30 | -82 | -8 |
| **R inf. Temporal** | 0.0001* | 1 | 5.65 | 54 | -49 | -26 |
| **R fusiform** | 0.0039* | 1 | 5.52 | 33 | -73 | -17 |
| **R cerebellum** | 0.0039* | 1 | 5.27 | 36 | -58 | -38 |
| **L fusiform** | 0.0039* | 2 | 5.26 | -36 | -64 | -11 |
| **R fusiform** | 0.0117* | 1 | 5.12 | 33 | -76 | -8 |

*FWE corrected p-value. Cluster sizes are reported at the p = 0.0039 (permutations).

Thereafter, as described before, we analyzed the non-face specific contrast at the coordinates x = 16, y = 13, z = 15 vxls. The most significant peak was x = 30, y = - 79 and z = - 8 mm, in the fusiform area (⬄ x = 17, y = 12, z= 15 vxls; p = 0.0039, FWE correction), near the seed pattern. Moreover, among significant peaks, we found some of them in the contralateral fusiform area as previously in the SPM analysis (Table S4).

In summary, comparably to what we found with SPM analysis, the results found with SnPM showed that significant correlations with the seed pattern can be obtained for both face-specific and non-face-specific contrasts.

**Table S4 Coordinates of significant correlations and their anatomic location. (Visual vs Random. Seed: right fusiform. x = 33, y = -76, z = -8 mm .Contrast: non-face specific)**

| **Anatomic location** | **p (corr) Voxel level** | **Cluster size (K)** | **Pseudo t** | **x** | **y** | **z** |
| --- | --- | --- | --- | --- | --- | --- |
| **R fusiform** | 0.0039* | 325 | 11.65 | 30 | -79 | -8 |
| **R cerebellum** | 0.0039* |  | 7.75 | 36 | -73 | -26 |
| **L cerebellum** | 0.0039* |  | 7.38 | -33 | -67 | -29 |
| **L fusiform** | 0.0039* |  | 7.07 | -21 | -82 | -8 |
| **R MCC (Middle Cingulate Cortex)** | 0.0039* | 6 | 7.25 | 12 | 17 | 40 |
| **L MCC** | 0.0039* |  | 5.95 | -9 | -7 | 46 |

*FWE corrected p-value. Cluster sizes are reported at the p = 0.0039 (permutations). Cognitive interpretations are limited for this contrast (non-specific visual activations).

##### Cross-correlation analysis in the auditory area

Next, in order to test whether the analysis reflects spatial information, which is specific to cognitive processing, as previously in SPM, we used the seed pattern for the face-specific contrast at the coordinates x = 10, y = 32, z = 21 voxels (⬄x = 51, y = -19, z = 10 mm) in the auditory cortex. The most significant correlation peak was in x = 51, y = - 19 and z = 10 mm which corresponds to the seed pattern coordinates (Table S5).

**Table S5 Coordinates of significant correlations and their anatomic location. (Auditory vs. Random. Seed: Right auditory area. x = 51, y = -19, z = 10 mm. Contrast: face-specific)**

| **Anatomic location** | **p (corr) Voxel level** | **Cluster size (K)** | **Pseudo t** | **x** | **y** | **z** |
| --- | --- | --- | --- | --- | --- | --- |
| **R primary auditory area (Heschl)** | 0.0039* | 53 | 13.83 | 51 | -19 | 10 |
| **R sup temporal** | 0.0195* | 2 | 5.09 | 66 | -28 | 7 |
| **R SupraMarginal** | 0.0195* | 1 | 5.16 | 60 | -43 | 37 |
| **R inf parietal lobule** | 0.0391* | 1 | 4.98 | 54 | -43 | 46 |

*FWE corrected p-value. Cluster sizes are reported at the p = 0.0039 (permutations).

Similarly to what we did with SPM, we also compared the results of using a seed pattern in the fusiform area with the results of a seed in the auditory region. The results of this analysis (Table S6) were similar to those obtained using randomized images as the statistical baseline.

**Table S6 Coordinates of significant correlations and their anatomic location. (Face vs. Auditory. Seed: Right fusiform. x = 33, y = - 76, z = - 8 mm. Contrast: face-specific)**

| **Anatomic location** | **p (corr) Voxel level** | **Cluster size (K)** | **Pseudo t** | **x** | **y** | **z** |
| --- | --- | --- | --- | --- | --- | --- |
| **R fusiform** | 0.0039* | 453 | 15.91 | 33 | -76 | -8 |
| **R cerebellum** | 0.0039* |  | 7.70 | 51 | -52 | -35 |
| **L fusiform** | 0.0039* | 25 | 5.94 | -24 | -82 | -11 |
| **L orbitofrontal** | 0.0039* | 53 | 5.53 | -12 | 50 | -8 |
| **L cerebellum** | 0.0313* | 7 | 4.82 | -48 | -55 | -32 |

*FWE corrected p-value. Cluster sizes are reported at the p = 0.0039 (permutations).

### Supplementary Methods

#### Analysis with SPM

In order to perform the cross-correlation of patterns of brain activity in a given contrast image, the files “cross.exe”, “filelist.txt” and “cmd.exe” and the initial contrast image are put in the same folder. Thereafter, we open the “cmd.exe” file and enter the command *cross* followed by the coordinates (x, y, z in voxels) of a chosen point near the peak of the activity of the pattern of interest (seed) and the radius of the pattern at the end (example: cross 45 5 12 5). The latter example means: find the cross-correlation of a pattern located in the position x=45, y=5, z=12 with a radius of 5 voxels. Before running the cross-correlation, one should indicate the file of interest (which will be analyzed) by indicating the path in “filelist.txt” (ex: D:\PROJECT\Sub02\Cross_corr\con_0006.img. During the analysis, the first check is at the given coordinates, where the correlation should be 1. If the initial correlation is 1, rotations step by step of the pattern are launched.

After running the cross correlation, we obtain the following NIFTI files: crosscorr_final.nii, crosscorr_final_Negative.nii, largeimage.nii (the NIFTI image with added margins), pattern.nii and the files indicating the angles of the rotated pattern (X_angles.nii, Xneg_angles.nii, Y_angles.nii, Yneg_angles.nii, Z_angles.nii, Zneg_angles.nii). The file “crosscorr_final.nii” contains all the results of the positive crosscorrelations and “crosscorr_final_Negative.nii” contains the negative ones. The “X_angles.nii”… indicate all the rotation angles of the pattern corresponding to each cross-correlation. The file “pattern.nii” contains the initial pattern of the activity that was compared to the whole brain activity. All these files are created by the analysis in the same folder with the initial NIFTI image.
